# GAPPadder: A Sensitive Approach for Closing Gaps on Draft Genomes with Short Sequence Reads

**DOI:** 10.1101/125534

**Authors:** Chong Chu, Xin Li, Yufeng Wu

## Abstract

**Background:** Closing gaps in draft genomes is an important post processing step in genome assembly. It leads to more complete genomes, which benefits downstream genome analysis such as annotation and genotyping. Several tools have been developed for gap closing. However, these tools don’t fully utilize the information contained in the sequence data. For example, while it is known that many gaps are caused by genomic repeats, existing tools often ignore many sequence reads that originate from a repeat-related gap.

**Results:** In this paper, we propose a new approach called GAPPadder for gap closing. The main advantage of GAPPadder is that it uses more information in sequence data for gap closing. In particular, GAPPadder finds and uses reads that originate from repeate-related gaps. We show that these repeat-associated reads are useful for gap closing, even though they are ignored by all existing tools. Other main features of GAPPadder include utilizing the information in sequence reads with different insert sizes and performing two-stage local assembly of gap sequences. We compare GAPPadder with GapCloser, GapFiller and Sealer on one bacterial genome, human chromosome 14 and the human whole genome with paired-end and mate-paired reads with both short and long insert sizes. Empirical results show that GAPPadder can close more gaps than these existing tools. Besides closing gaps on draft genomes assembled only from short sequence reads, GAPPadder can also be used to close gaps for draft genomes assembled with long reads. We show GAPPadder can close gaps on the bed bug genome and the Asian sea bass genome that are assembled partially and fully with long reads respectively. We also show GAPPadder is efficient in both time and memory usage. The software tool, GAPPadder, is available for download at https://github.com/Reedwarbler/GAPPadder.

## 1 Introduction

With the fast developing high-throughput sequencing technologies, de novo genome assembly from sequence reads has become a major application of sequencing technologies. So far many genome assembly software tools have been developed, including e.g. [3, 11, 18, 22]. As sequence data from many species is becoming increasingly more available, draft genomes of many species have been assembled. Furthermore, more recent sequencing technologies such as long reads sequencing are expected to lead to even more assembled genomes with better quality than before.

Despite all these exciting developments, it is still challenging to obtain complete genomes with the current technologies and assembly tools, especially at regions that are highly repetitive or have low coverage. At present, most assembled genomes contain gaps. For relatively complex genomes, only draft genomes which usually contain a large number of gaps are available. A more complete genome is highly desirable since it leads to better annotation, less genotyping error and easier identification of causal variation associated with traits [4] than a genome with many gaps. For example, 45 new avian species have been sequenced and assembled recently in a comparative study of avian genomes [21]. Draft genomes of 25 out of these 45 species have average N50 around 48 kb, which indicates the draft genomes are fragmented with many gaps. About 3,000 genes are likely missing or only partially annotated due to gaps. As a result, only 70% to 80% of the entire catalog of avian genes can be predicted, which may cause bias in downstream analysis.

With the development of the third generation sequencing technology, long reads from different platforms, like Pacific Biosciences, Illumina TruSeq, Oxford Nanopore, have been developed. With the help of these new technologies, the quality of the assembled draft genomes is greatly improved. In general, long reads are used in two ways to help to improve the draft genome assembly: 1) Long reads are used to scaffold the contigs and fill the gaps on the draft genomes assembled from high coverage short reads. 2) Long reads are directly used to assemble the draft genomes. Due to the high error rates of long reads, read depth is required to be high to guarantee the quality of genomes assembled directly from long reads, and thus sequencing cost can be high. In comparison, for scaffolding contigs and closing gaps with long reads, the coverage is usually not required to be very high. However, there are still gaps on the draft genomes even assembled with long reads, especially for draft genomes initially assembled high coverage short reads and then improved with long reads. Thus, it is still needed to close the gaps on draft genomes assembled with long reads. At present, short sequence reads are still the most available sequence reads. Thus, it is important to develop methods that can close gaps on draft genomes with short sequence reads that are readily available.

Several tools have been developed for closing gaps on draft genomes with short reads. GapCloser is a stand-alone tool in the SOAPdenovo [12] package. It performs several iterations of base extension steps using the reads aligned to specific regions. GapFiller [2] implements a method that finds read pairs with one end aligned within a contig and its mate partially aligned to the draft genome and partially located in a region identified as a gap. These partially aligned reads are used to close the gap through sequence overlapping. Sealer [16] generates pseudo long reads from paired-end sequence reads by filling the unknown sequences between read pairs using the redundancy in sequence coverage, and then the pseudo long reads are used to fill the gaps. While these approaches have been used to close gaps in assembled genomes, these tools still cannot close many gaps (especially those originated in more complex genomic regions, e.g. repeats).

In this paper, we develop a new approach called GAPPadder for closing gaps on draft genomes. Similar to tools such as GapCloser and GapFiller, GAPPadder also performs local assembly from reads that originate from gap regions. The following are the main features of GAPPadder and also differences between GAPPadder and the existing methods.

- GAPPadder uses more information about the gaps contained in sequence reads than existing methods. GAPPadder collects more reads relevant for gap closing, especially repeat-associated reads which are ignored by all the existing tools. Moreover, GAPPadder collects higher quality reads by utilizing more information with different insert sizes of pair-end (PM) and mate-pair (MP) reads.
- GAPPadder uses a different local assembly method for gap closing compared with existing methods. Existing methods often rely on local extension of contigs. GAPPadder, instead, performs a two-stage local assembly: it first assembles contigs in the gap and then generates higher quality local assembly of gap sequences by merging contigs.

We compare GAPPadder with existing approaches using real sequence data from staphylococcus aureus, human chromosome 14 from GAGE [17], and whole genome sequencing data (with PE and MP reads) of one human individual NA12878 from Illumina. These genomes are assembled from short reads only. We show GAPPadder can close more gaps than GapCloser, GapFiller and Sealer with these short sequence reads. Besides these draft genomes assembled with only from short reads, we also compare GAPPadder with GapCloser on two draft genomes assembled with long reads: the bed bug draft genome assembled with hybrid short and long reads and the Asian sea bass draft genome directly assembled from long reads. We show many gaps can be fully closed and extended by GAPPadder and GapCloser, and GAPPadder closes much more than GapCloser on the hybrid assembled bed bug genome.

### 1.1 Gaps in draft genomes

De novo assembly of reads produces contigs. Contigs are then further linked with paired-end (PE) or mate-pair (MP) reads to form scaffolds. Scaffolds contain multiple gaps, whose lengths are estimated from the insert sizes of PE or MP reads. In general, extension of contigs stops at sites with repetitive regions, heterozygous alleles, sequencing errors or low read coverage [14, 20]. Gaps can be mainly classified to three types. The most common type is the repeat-associated gap. Repeat is a piece of DNA which may have multiple copies in the genome. Note that these copies may differ slightly from each other. There are different types of repeats, including LINE, SINE, LTR elements, DNA transposon, satellites, etc. Repeat-associated gaps can be categorized to be satellite-associated, dispersed low divergent repeats-associated, and tandem repeats-associated. We show the results of masking the gap regions on chromosome 14 of human using RepeatMasker [19] in Fig. 1. To get the gap regions of the draft genome of chromosome 14, which is assembled by ALLPATH-LG and released in GAGE, we align the flanking regions to the reference genome, and thus get the benchmarked gap sequences (i.e. sequences from the reference genome that are missing in the draft genome). One can see that over 90% of the gaps are masked as repeat-associated gaps. Therefore, to develop gap closing methods, it can be very useful to consider the implications of repeats on gaps.

**Figure 1.**
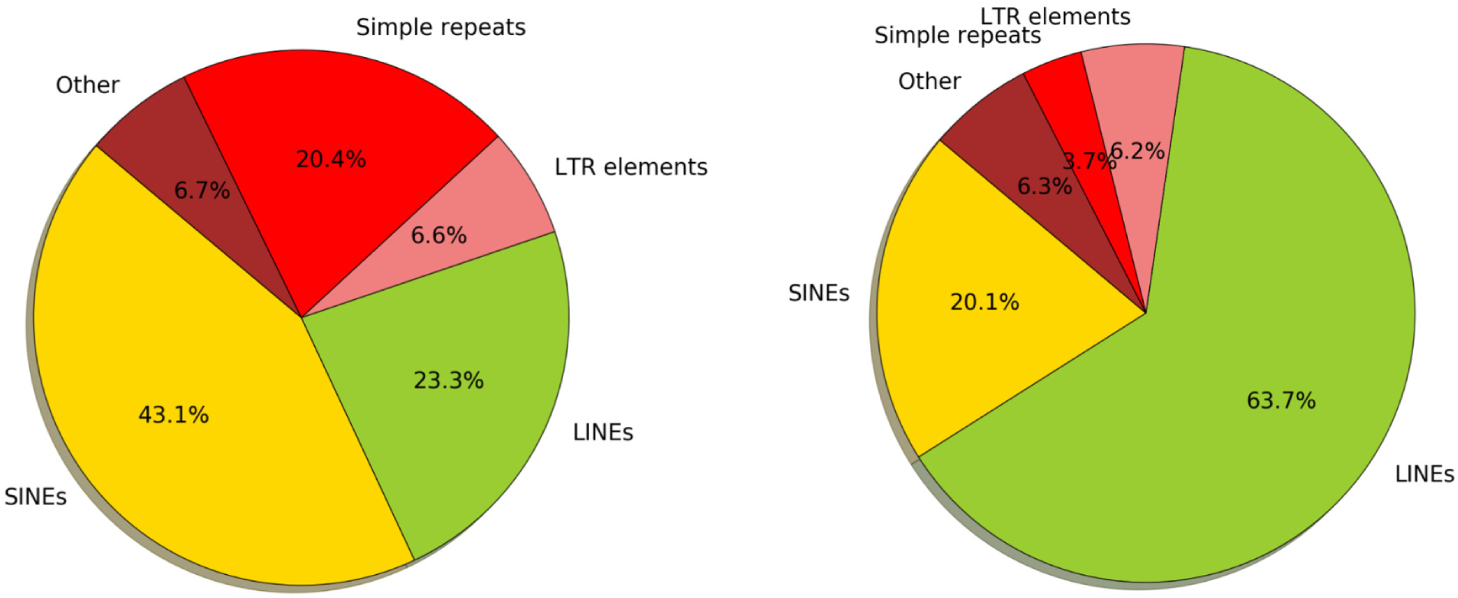
Percentage of the masked gap sequences of each type of repeats. 3,934 gaps are extracted from the draft genome of human chromosome 14 that is released in GAGE. By aligning the flanking sequences to the human reference, we extract the gap sequences. We use RepeatMasker to get the types of repeats (e.g. LINE, SINE, LTR elements, etc) of these gap sequences. One gap may be masked to multiple repeat types. The left part shows the percentage of masked gaps for each repeat type. The right part shows the percentage of masked bases for each repeat type.

**Figure 2.**
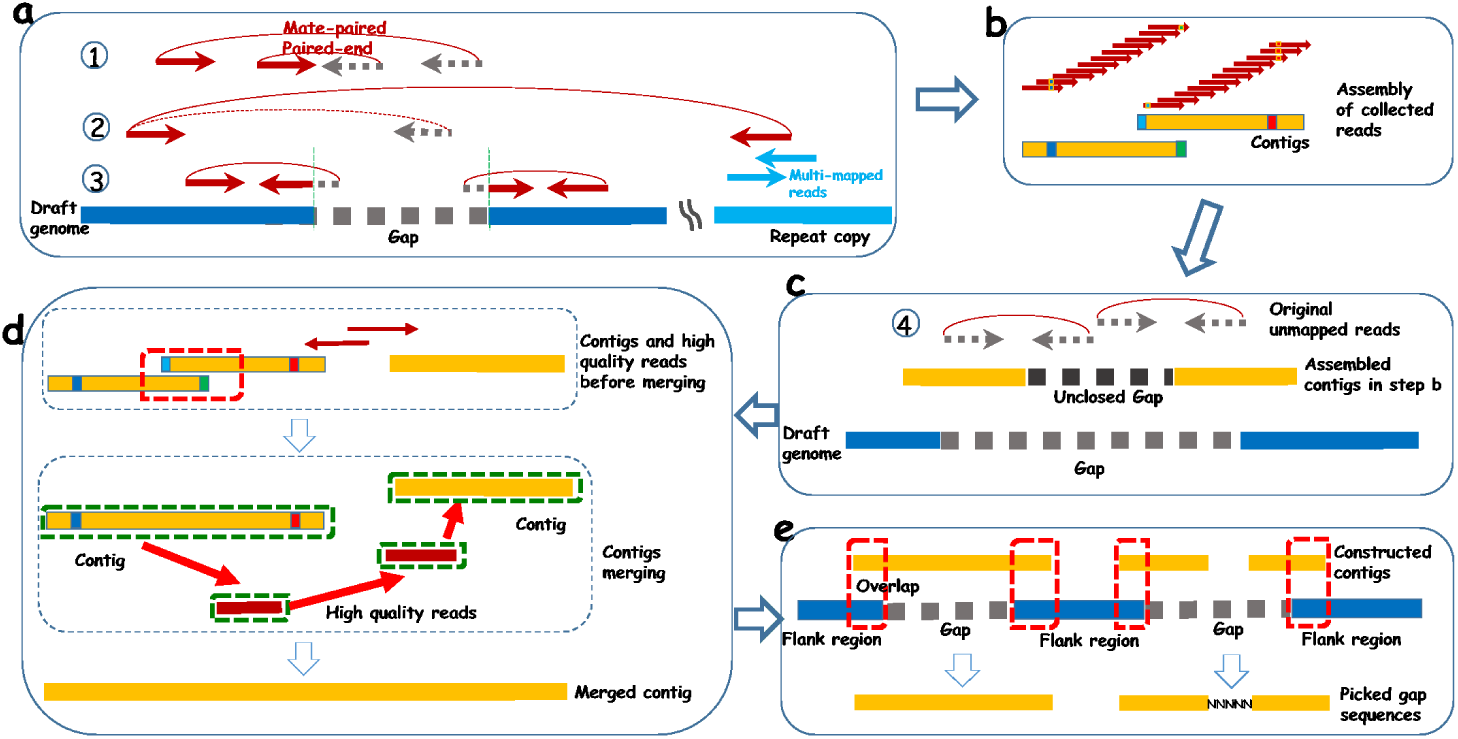
**Pipeline of GAPPadder for closing gaps:** (a) Align reads back to the draft genome and collect the first three types of relevant reads originated from the gaps. (b) Perform local assembly with the collected reads, and merge contigs overlapping with each other to generate more complete gap sequences. (c) Align the unmapped reads to the constructed contigs of each gap, and collect the aligned (also their mate) reads. (d) Merge the contigs to form more complete assembly. (e) Gap sequences are obtained by aligning the merged contigs to the flanking regions of the gaps.

## 2 Results

We compare GAPPadder with GapCloser, GapFiller and Sealor on datasets of three draft genomes of different sizes and with known reference sequences: staphylococcus aureus, human chromosome 14 and human whole genome. Data of staphylococcus aureus and human chromosome 14 are from GAGE [17]. We choose the draft genome assembled by ALLPATH-LG. For staphylococcus aureus, two groups of high coverage data of different insert sizes are used. While for the human chromosome 14, three groups of data of different insert sizes are used. The data with long jump library is of very low coverage. The human whole genome (NA12878) high-coverage PE and MP sequence reads are from Illumina. The draft genome of NA12878 is released in [7], which is assembled by ALLPATH-LG. Detailed information of the four datasets are given in the additional file [see Additional file 1].

### 2.1 Comparison with existing tools

In Table 1, we show the results of GAPPadder and the other three tools on staphylococcus aureus, human chromosome 14 and human whole genome. Note that Sealer only runs well on short insert size data. For data with very long insert size, it can be extremely slow. So when running Sealer on the human whole genome data, we do not use the long insert size data. Detailed commands and parameters of running each tool are provided in the additional file [see Additional file 1]. The results show that GAPPadder outperforms the other three tools on the three datasets. For S. aureus and H. chromosome 14 datasets, GAPPadder closes more gaps than the other three tools. For the human whole genome datasets, GapCloser runs out of memory (on a server with 256G memory) and GapFiller did not finish after running for more than 725 hours. In comparison, GAPPadder and Sealer respectively close 130,371 and 110,876 gaps out of the 220,318 gaps.

**Table 1.**
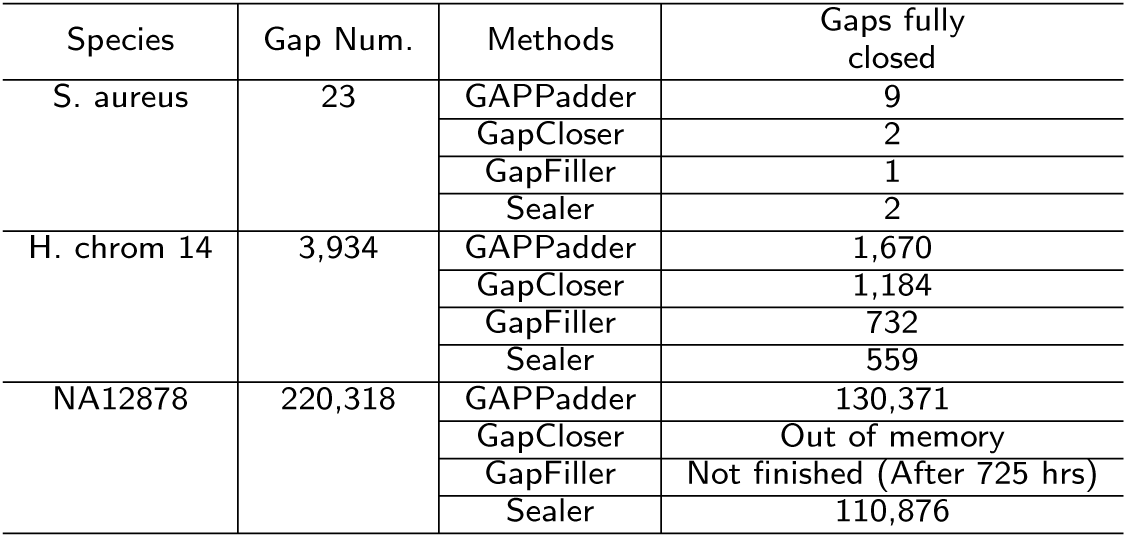
Comparison of the four tools on three datasets: S. aureus, human chromosome 14 and whole human genome for NA12878, whose draft genomes have 23, 3,934, and 220,318 gaps respectively. Overall, GAPPadder closes more gaps than the other three tools on the these datasets.

To show the effect of the repeat-associated reads, we run a revised version of GAPPadder that does not use these repeat-associated reads for gap filling. This “streamlined” version of GAPPadder closes 1,103 gaps, much less than the original version of GAPPadder, which closes 1,670 gaps. This indicates that repeat-associated reads are indeed useful for gap closing.

In Fig. 3, we compare the four tools on different ranges of gap lengths of the closed gaps. The left part shows the distribution of gap length of all the 3,934 gaps on the draft genome of human chromosome 14. Over 80% of the gaps are shorter than 1k, and over 95% of the gaps are smaller than 2k. The right part shows the number of fully closed gaps of the four tools on different ranges of gap length. GAPPadder significantly outperforms the other three tools on gaps shorter than 1 kb, while GAPCloser performs slightly better on gaps longer than 1 kb.

**Figure 3.**
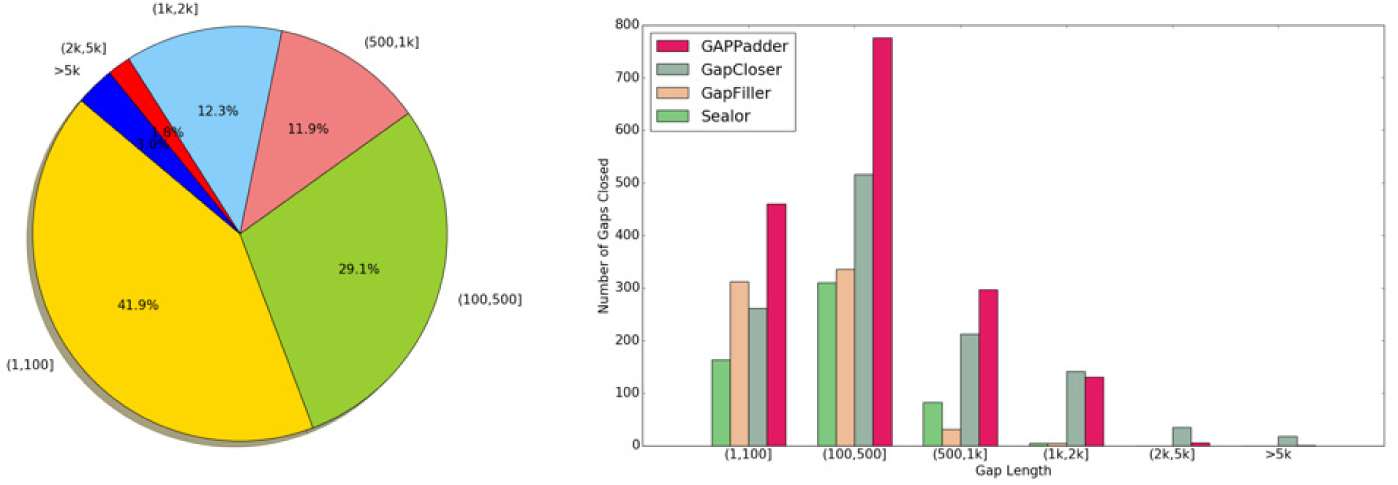
**Comparison on different gap length:** The left part shows the distribution of gap length of all the 3,934 gaps on the draft genome of human chromosome 14. Over 80% of the gaps are smaller than 1 kb, and over 95% of the gaps are smaller than 2 kb. The right part shows the number of fully closed gaps of the four tools on different ranges of gap length.

### 2.2 Comparison on data with different insert sizes

For assembling draft genomes, usually data of different insert sizes are provided. Paired-end reads of long insert size or mate-paired reads can be helpful for closing (especially long) gaps on the draft genomes. Because of the different strategies used, the performance of different tools differs significantly on datasets of different insert sizes. To evaluate the performance of the four tools on different insert size datasets, we compare the four tools on the human chromosome 14 datasets with only short insert size data, only long insert size data, and combined data with short and long insert size. The results are shown In Table 2. With data with short insert sizes, GapCloser performs the best. But with only long inset size data, GAPPadder significantly outperforms the other three tools. For comparison on the combined dataset with reads of both short and long insert sizes, GAPPadder performs the best.

**Table 2.** Comparison of the four tools on different insert size data. Three groups of data of insert sizes (180, about 2,500 and about 35 kb) of human chromosome 14, and their combination are used for comparison. Results are given for reads with 180bp insert size only, and reads with 2,500bp insert size only and combined reads (with 180bp, 2,500bp and 35kb insert sizes).

| Methods | Insert size |  |  |
| --- | --- | --- | --- |
|  | 180 | 2,283 to 2,803 | Combined |
| GAPPadder | 862 | 1,481 | 1,670 |
| GapCloser | 1,142 | 216 | 1,184 |
| GapFiller | 484 | 173 | 732 |
| Sealer | 468 | 308 | 559 |

### 2.3 Time and memory usage

All four tools are benchmarked on a 64-core server with AMD 6380 CPU @2.499 GHz and 256 GB RAM. To compare the time and memory usage of these four tools, we benchmark the four tools on the human chromosome 14 datasets. When running Sealer, we set the maximum allowed memory to 40G, and other parameters are set as suggested by its manual. For GapCloser, we use the default parameters. For GapFiller, the parameter for the number of iterations to run is set to be 5. See the additional file [Additional file 1] for more detailed information of running the tools. In terms of running time, GapCloser, GapFiller, Sealer and GAPPadder take 30m 32s, 424m 47s, 160m 23s, and 85m 12s respectively. For memory usage, GapCloser takes 7.8G at the peak, Sealer takes 40G (as set in the parameter), while GapFiller and GAPPadder take less than 2G memory. Therefore, GapCloser is the most efficient one among the four tools, but it requires more memory. GAPPadder is slightly slower than GapCloser but uses much less memory.

### 2.4 Closing gaps on draft genomes assembled partially or fully with long reads

Although long reads help to improve the draft genome assembly, large number of gaps may still remain in the draft genome, especially for the draft genome originally assembled from short reads and then improved from long reads. To evaluate the performance of GAPPadder on two draft genomes that are partially and fully assembled with long reads, we run GAPPadder on two draft genomes: 1) The bed bug cimex lectularius draft genome (released in [24]) which is assembled with hybrid data of both short and long reads, 2) Asian sea bass draft genome (released in [23]) that is purely assembled from high coverage PacBio long reads. The bed bug genome is initially assembled with 73x coverage Illumina short reads using ALLPATHS-LG [3] assembler. And then Illumina Moleculo kit is used to sequence long reads with average length 3,500bp, which is used to improve the initial assembled draft genome. However, even for the improved draft genome, there are still many gaps. In the final released assembled genome, there are 118,821 gaps, out of which 97,251 gaps are larger than 100bp. We run GAPPadder and GapCloser to close the gaps. As some gaps are really small (just several bases), and to evaluate the power of different tools we only focus on these 97,251 gaps that are larger than 100bp. Three sets of Illumina short reads with insert sizes of 185, 367, and 3,000bp and coverage of 34x, 12x, and 7x respectively are used for gap closing. GAPPadder reports 19,476 gaps are fully closed and 52,879 gaps are partially extended, while GapCloser reports 3,299 and 2,417 are fully closed and partially extended respectively. To validate the fully closed and partially extended gaps, for each closed gap sequence, we extract the left and right flank regions of length 150bp each, and concatenate them with the gap sequence. Then we align the reads back to the concatenated sequences and check whether there are reads clipped at the joint regions. If enough (by default 10) reads are fully mapped at the joint regions and over 95% of the bases (for extended ones, excluding the not-filled regions) of each sequence at least have 10 reads covered, then we call this gap is validated. Otherwise it is not. For GAPPadder, 14,925 fully closed gaps and 37,802 partially extended gaps are validated in this way. While for GapCloser, 2,737 fully closed and 20 partially extended gaps are validated. In table 3 we show the comparison.

**Table 3.** Evaluation of GAPPadder and GapCloser on closing gaps for bed bug draft genome. The draft genome is initially assembled with high coverage short reads, and then improved with long reads. GAPPadder fully closes 14,925 out of reported 19,476 gaps and extends 37,802 out of reported 52,879 gaps. As a comparison, GapCloser fully closes 2,737 (3,299 are reported) gaps and extends 20 (2,417 are reported) gaps.

| Category |  | GAPPadder | GapCloser |
| --- | --- | --- | --- |
| Fully closed | Reported | 19,476 | 3,299 |
|  | Validated | 14,925 | 2,737 |
| Extended | Reported | 52,879 | 2,417 |
|  | Validated | 37,802 | 20 |

For the Asian sea bass draft genome, it is primarily assembled from 90x PacBio data and then scaffolded using transcriptome data. From the release draft genome, 110 gaps are extracted and all of them are larger than 100bp. We run GAPPadder and GapCloser to close the gaps. Two sets of Illumina short paired end reads with insert sizes of 500bp and 750bp, read length 100bp, and total coverage 80x are used for closing the gaps on the draft genome. For GAPPadder, 14 and 47 gaps are reported to be fully closed and partially extended respectively. We use the same validation approach as used in validating the gap sequences of the bed bug genome, and 5 fully closed and 13 partially extended gaps are validated in this way. For GapCloser, 46 and 41 are reported to be fully closed and partially extended respectively, and 6 and 1 out of the fully closed and partially extended gaps are validated by the the same way.

## 3 Discussion and Conclusions

In this paper, we propose a sensitive approach for closing gaps on draft genomes with paired-end reads and mate-paired reads. Empirical results show that when both short and long insert size data are provided, our tool GAPPadder outperforms GapCloser, GapFiller and Sealer. This is likely due to the fact that GAPPadder uses more reads (especially the repeat-associated reads) to close the gaps which are ignored by all other tools. Besides that, GAPPadder takes advantage of long insert size data and performs a two-stage local assembly approach to construct more complete gap sequences. In Figure 4, we show the comparison of the four tools on closing one example gap, which is about 770bp long on chromosome 14. GapCloser only extends a little on the left part. GapFiller and Sealer even have no extension at all, and thus are not shown in the UCSC Genome Browser. In comparison, GAPPadder fully closes the gap. One possible reason is the gap is composed by part of a SINE copy and part of a LINE copy as shown in the UCSC genome browser. The repeat-associated reads used by GAPPadder provide enough coverage for assembling the gap region.

**Figure 4.**
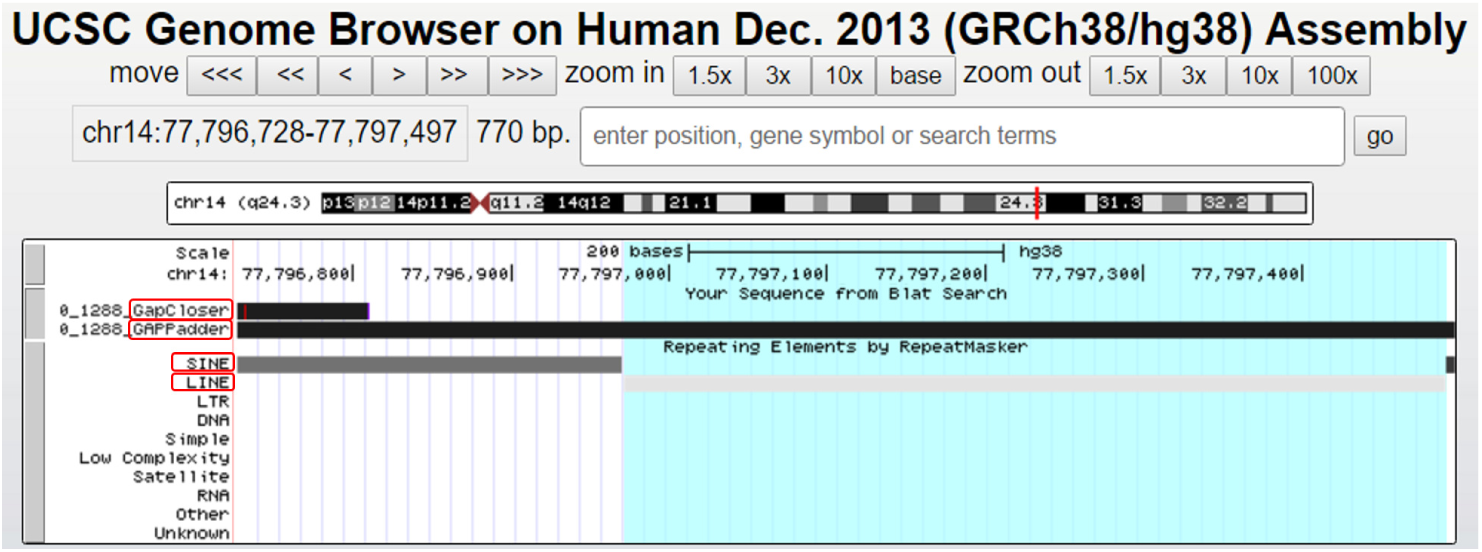
One example gap closed by GAPPadder but failed by other three tools: one gap on the chromosome 14 draft genome is fully closed by GAPPadder, but other three tools fail to fully close it. The gap is about 770bp long, and from the UCSC genome browser we can see it is composed by part of a SINE copy and part of a LINE copy. GapCloser only extends by a short length on the left part. GapFiller and Sealer fail to fill this gap.

However, when only using reads with short insert size for closing the gaps on human chromosome 14, GAPPadder does not performs as well as GapCloser. One reason is that GAPPadder relies on the contigs constructed in the first step to collect both unmapped reads. If the insert size is small, then the collected reads mainly come from the two ends of the gap, and thus the middle part will be difficult to construct in the following steps. In comparison, GapCloser uses an iterative strategy which can gradually extend the contigs. This indicates that tools are designed with different strategies, and users should choose a tool based on the kind of data.

For draft genomes directly assembled from high coverage long reads, often the draft genomes contain far less gaps than those assembled from short reads. One reason is for less complex genomes, chromosome level contigs are directly assembled which do not need to do scaffold and gap closing. Second, very little scaffolding tools are developed for these near completed draft genomes, thus even gaps exists, they are not reported in the released draft genomes. Nonetheless, we observe that there can still be gaps within draft genomes that are directly assembled from long reads. Our results indicate that our GAPPadder tool can still be useful in the age of long reads genome assembly.

One possible future research on gap filling is incorporating long reads to close the gaps on the draft genomes. Direct assembly of long reads usually requires the coverage should be high enough to get a high quality draft genome, which usually leads to high sequencing cost. Although low coverage of long reads cannot provides a high quality draft genome, it may help to close the gaps on the draft genome generated from short reads, especially for the long duplicate-associated or repeat-associated gaps.

## 4 Methods

### 4.1 High level approach

In this paper, we propose GAPPadder for closing gaps on draft genomes, which greatly improves the sensitivity. The key idea is that GAPPadder utilizes more information (i.e. relevant reads originated from the gaps) contained in the sequence reads for gap filling. For example, GAPPadder collects the repeat-associated reads, which are ignored by all existing approaches. Our main observation is that reads originated from repeat-associated gaps may be mapped to other copies of the same repeat contained in the genome. Therefore, the two ends of these read pairs may be discordantly mapped (i.e. mapping positions of the two ends are much farther away from each other than expected on the same chromosome or even located at different chromosomes). GAPPadder also uses multi-mapped reads near these reads because they may also be useful for the assembly of gap sequences, especially when the collected reads are of low coverage. GAPPadder utilizes the long insert size reads or mate-pair (MP) reads to collect high quality reads. Another important step in GAPPadder is that it performs two-stage local assembly for each gap: it first assembles contigs from relevant reads in the gap; then it merges these contigs to construct long gap sequences. The main observation is that assembled gap sequences usually are in the form of relatively short segments (contigs) due to positions with errors or variations. These contigs overlap but are usually not assembled by standard assembly methods into longer sequences due to mismatches between contigs. The merging step implemented in GAPPadder allows the merging of these contigs to form long (sometimes complete) gap sequences.

### 4.2 Relevant reads originated from gap regions

Similar to several existing methods, GAPPadder starts by finding relevant reads that originate within each gap. In this paper, we are mainly concerned with paired-end (PE) or mate-paired (MP) reads. When aligning the reads back to the draft genome using tools e.g. BWA [10], four types of read pairs can be considered to originate from the gap regions. All these read pairs are located near the gap under consideration. This is shown in Figure 2.

i. One end mapped and its mate unmapped. For a read pair, suppose the left (respectively right) read is aligned (by default with mapping quality greater than 30), and the alignment position is within *m* + 3*υ* distance from the left (respectively right) breakpoint of the gap. Then this read is called the anchored read. Here *m* and *υ* are the mean and standard derivation of insert size respectively. Further suppose the mate of the anchored read is unmapped. Then the unmapped read comes from the gap region with high probability.
ii. Discordant reads caused by repeats or duplicate segments. If one read of a pair comes from the gap region, then when aligning the read back to the draft genome, this read will be unmapped. However, if the gap region comes from a repeat region and there are other copies of the repeat that are already included in the draft genome, then this read may be aligned to another repeat copy. As a result, both ends of the pair will be mapped, but become discordant (with insert size outside the range [*m* − 3*υ, m* + 3*υ*]) or are mapped to different chromosomes. This kind of reads may originate within the gap and may help the assembly of gap sequences. Besides the discordant reads, multi-mapped reads (by default with mapping quality 0) near the discordant reads are also useful for assembly. This is because if the gap is repeat-associated, these multi-mapped reads from the copy of the same repeat can be useful, especially when collected reads have low coverage.
iii. Reads clipped at the breakpoints of the gaps. For the reads overlapping the breakpoints, parts of the reads will be aligned to the draft genome, and the other parts will be clipped. Clipped reads are useful to extend the assembled regions from collected reads to both sides of the flanking regions of gaps. This allows the assembled gap sequences to be positioned in the draft genome.
iv. Both reads of a pair are unmapped reads. When the gap is long enough, then both ends of a pair likely originate within the gap region. As a result, when aligning reads back to the draft genome, both reads will be unmapped. Such unmapped reads may play an important role if the insert size is short and the gap is long. In this situation, it is difficult to find anchored reads. As a result, the middle part of the gap will not be filled using reads with anchor. We note that unmapped reads may be just due to reads errors and thus irrelevant for gap filling. The challenge is that we do not know which unmapped reads indeed originate from some gap, and if so, which gap they originate. We will explain how to address this problem in the following sections.

Most existing tools use only the type-iii reads, while GAPPadder uses all four types of reads.

### 4.3 Gap closing procedure

As shown in Fig. 2, there are five steps of GAPPadder. We process each gap in the draft genome independently. First, we collect the first three types of reads that may originate from a gap. Second, we perform local assembly of the collected reads of each gap. This generates (usually short) contigs that are segments of the gap sequences. Then, we align the unmapped reads to the constructed contigs and collect the aligned (also their mate) reads. We merge the contigs to form more complete assembly using a customized designed algorithm. Here, the high quality reads are treated as short contigs and is used for contig merging. Finally, we fill the gaps by aligning the merged contigs to the flanking regions of the gaps.

#### 4.3.1 Collection of gap-associated reads

GAPPadder allows PE or MP reads of different insert sizes. For each group of reads of one specific insert size, we collect reads separately and then all these reads are used together for gap closing. To collect reads for one specific insert size, we first align the reads back to the draft genome using BWA.

We search for type-i reads that are mapped within *m* + *3υ* + *l* (where *l* is the read length) distance from the breakpoints, and their mate reads are unmapped. The mapped reads are used as anchor, and the unmapped mate reads are used for gap closing. Here we consider all possible anchor reads, even when their mapping quality scores are low.

For type-ii reads, we search for reads in the region [*b*_1_ − *m* − 3*υ* − *l, b*_2_ + *m* + 3*υ* +*l*], where *b*_1_ and *b*_2_ are the breakpoint positions of the gap. If a read A falls in this region but its mate read B is aligned outside the region, and also the mapping quality of read B is 0, then read B is considered to be type-ii. Also, suppose read B is aligned at position p, then we also use the reads whose mapping quality is 0 and aligned within the region [*p* − *d/2, p*+*d*/2], where d is the gap length. This is because a read with mapping quality 0 is with high probability to be a multi-mapped read.

For type-iii reads, the assembly quality at the end of contigs is usually low. Thus, when collecting reads clipped at breakpoints, we set some slack value (by default 20bp) to allow some distance between the clip position and the breakpoints of the gaps. Note that one read may satisfy the conditions of more than one gaps. And if this happens, we let the read to be used for all the related gaps.

Out of these collected reads, we define those reads whose mates (anchor reads) are uniquely mapped as high quality reads. Here, if the mapping quality of a read is equal to 60, then the read is considered to be uniquely mapped. In other words, we believe that with high probability these reads are from the specific gap region. We also collect the unmapped reads which will be used in the third step.

#### 4.3.2 Local assembly of collected reads

This is the first stage of our two-stage local assembly approach. Once the reads are collected, we perform local assembly with the reads of each gap. KMC2 [6] is used to convert the reads to k-mers, then Velvet [22] is used to assemble the kmers to contigs. This step is similar to the repeat assembly approach developed in Chu *et al*. [5].

#### 4.3.3 Collection of type-iv reads with the constructed contigs

From the previous steps, we construct contigs for each gap from the collected reads. If the insert size is shorter than the gap length, then both reads of a read pair may be unmapped. Such unmapped reads can be useful to construct longer contigs. This is still important even there are both paired-end and mate-pair reads of different insert sizes, and the insert sizes are longer than the gap sizes. Mainly because the coverage of mate-pair reads is usually low. As usually the mate-pair reads are initially used for scaffolding, but not for gap filling, and thus the coverage is usually not high, because of which the regions will still be constructed to pieces. So it is quite necessary to collect the both unmapped reads.

The challenge here is that we do not know which read pair comes from which gap, since they are unmapped. To solve this problem, we first collect all the unmapped reads. Then we align all the unmapped reads to the constructed contigs of each gap using BWA. By collecting the mapped reads, we collect the originally unmapped reads (now aligned to contigs of each gap) and their mate reads for each gap. Note that after the first-round assembly, we exclude those gaps that have been fully closed (see section 4.3.5 for details) from consideration. Then we only collect the unmapped reads for those not fully constructed.

#### 4.3.4 Merging contigs

This is the second stage of the two-stage local assembly approach. The previous steps often generate more than one contigs for each gap. In order to obtain a complete gap sequence, GAPPadder performs a contig merging step. Similar to the general genome assembly problem, contig merging can be performed based on prefix-suffix overlap between two contigs. We use the contig merging procedure in Chu *et al*. [5], which was originally developed for merging contigs for the repeat construction problem. Refer to Chu *et al*. [5] for more details on this procedure. As mentioned in section 4.1, for some regions of gaps, even though we have collected reads that fall into these regions, there may not be enough reads covering these regions. As a result, when we perform local assembly for these gaps, only short contigs (with little overlap with other contigs) are obtained for these regions, and usually they do not have overlap. A simple solution is that we can view these reads as contigs and include them in the contigs merging step. To improve the merging efficiency and accuracy, we only use the high quality (the mate reads are uniquely mapped) reads that cannot be aligned to the constructed contigs.

#### 4.3.5 Finishing gap sequence assembly

After contig merging, for each gap, there can be several constructed sequences. Most of these sequences are pieces of the repeats or wrongly assembled. So we need to identify the right one. We first check whether the whole gap is constructed. To identify the fully constructed ones, for each gap we get the two flanking sequences of the gap (by default 300bp for each). Then we align the two flanking sequences to the constructed contigs of the gap. If the left flanking sequence overlap with the left (right) side of the contig and the right (left) flanking sequence overlap with the right (left) side of the contig, and the two overlaps are of the same orientation (both are reverse complementary or both not), then we choose the contig as the gap sequence. If more than one contigs are found, we choose the longest one. In our experiments, we notice that for most of the filled gaps, there is usually only one satisfying these conditions. If complete gap sequences cannot be found, we choose the one that covers the gap the most.

## Author’s contributions

Conceived and designed the experiments: YW CC. Performed the experiments: CC. Analyzed the data: CC XL. Contributed reagents/materials/analysis tools: CC XL. Wrote the paper: YW CC XL.

## Acknowledgements

This research is supported in part by grants IIS-0953563 and IIS-1526415 from National Science Foundation. References

## Additional Files

Additional file 1 — Data and commands used in the experiments

Data used in the experiments, parameters and commands used for running the tools.

## References

1. Berlin, K., Koren, S., Chin, C.-S., Drake, J. P., Landolin, J. M., and Phillippy, A. M. (2015). Assembling large genomes with single-molecule sequencing and locality-sensitive hashing. Nature biotechnology, 33(6), 623–630.

2. Boetzer, M. and Pirovano, W. (2012). Toward almost closed genomes with gapfiller. Genome biology, 13(6), 1.

3. Butler, J., MacCallum, I., Kleber, M., Shlyakhter, I. A., Belmonte, M. K., Lander, E. S., Nusbaum, C., and Jaffe, D. B. (2008). Allpaths: de novo assembly of whole-genome shotgun microreads. Genome research, 18(5), 810–820.

4. Chaisson, M. J., Wilson, R. K., and Eichler, E. E. (2015). Genetic variation and the de novo assembly of human genomes. Nature Reviews Genetics.

5. Chu, C., Nielsen, R. and Wu, Y. (2016). REPdenovo: Inferring De Novo Repeat Motifs from Short Sequence Reads. PloS One, 11(3), e0150719.

6. Deorowicz, S., Kokot, M., Grabowski, S., and Debudaj-Grabysz, A. (2015). Kmc 2: Fast and resource-frugal k-mer counting. Bioinformatics, 31(10), 1569–1576.

7. Gnerre, S., MacCallum, I., Przybylski, D., Ribeiro, F. J., Burton, J. N., Walker, B. J., Sharpe, T., Hall, G., Shea, T. P., Sykes, S., et al. (2011). High-quality draft assemblies of mammalian genomes from massively parallel sequence data. Proceedings of the National Academy of Sciences, 108(4), 1513–1518.

8. Gordon, D., Huddleston, J., Chaisson, M. J., Hill, C. M., Kronenberg, Z. N., Munson, K. M., Malig, M., Raja, A., Fiddes, I., Hillier, L. W., et al. (2016). Long-read sequence assembly of the gorilla genome. Science, 352(6281), aae0344.

9. Kosugi, S., Hirakawa, H., and Tabata, S. (2015). Gmcloser: closing gaps in assemblies accurately with a likelihood-based selection of contig or long-read alignments. Bioinformatics, 31(23), 3733–3741.

10. Li, H. and Durbin, R. (2009). Fast and accurate short read alignment with burrows-wheeler transform. Bioinformatics, 25(14), 1754–1760.

11. Li, R., Zhu, H., Ruan, J., Qian, W., Fang, X., Shi, Z., Li, Y., Li, S., Shan, G., Kristiansen, K., et al. (2010). De novo assembly of human genomes with massively parallel short read sequencing. Genome research, 20(2), 265–272.

12. Luo, R., Liu, B., Xie, Y., Li, Z., Huang, W., Yuan, J., He, G., Chen, Y., Pan, Q., Liu, Y., et al. (2012). Soapdenovo2: an empirically improved memory-efficient short-read de novo assembler. GigaScience, 1(1), 1.

13. Manolio, T. A., Collins, F. S., Cox, N. J., Goldstein, D. B., Hindorff, L. A., Hunter, D. J., McCarthy, M. I., Ramos, E. M., Cardon, L. R., Chakravarti, A., et al. (2009). Finding the missing heritability of complex diseases. Nature, 461(7265), 747–753.

14. Miller, J. R., Koren, S., and Sutton, G. (2010). Assembly algorithms for next-generation sequencing data. Genomics, 95(6), 315–327.

15. Myers, E. W. (1995). Toward simplifying and accurately formulating fragment assembly. Journal of Computational Biology, 2(2), 275–290.

16. Paulino, D., Warren, R. L., Vandervalk, B. P., Raymond, A., Jackman, S. D., and Birol, I. (2015). Sealer: a scalable gap-closing application for finishing draft genomes. BMC bioinformatics, 16(1), 230.

17. Salzberg, S. L., Phillippy, A. M., Zimin, A., Puiu, D., Magoc, T., Koren, S., Treangen, T. J., Schatz, M. C., Delcher, A. L., Roberts, M., et al. (2012). Gage: A critical evaluation of genome assemblies and assembly algorithms. Genome research, 22(3), 557–567.

18. Simpson, J. T., Wong, K., Jackman, S. D., Schein, J. E., Jones, S. J., and Birol, I. (2009). Abyss: a parallel assembler for short read sequence data. Genome research, 19(6), 1117–1123.

19. Smit, A. F., Hubley, R., and Green, P. (1996). Repeatmasker open-3.0.

20. Treangen, T. J. and Salzberg, S. L. (2012). Repetitive dna and next-generation sequencing: computational challenges and solutions. Nature Reviews Genetics, 13(1), 36–46.

21. Zhang, G., Li, C., Li, Q., Li, B., Larkin, D. M., Lee, C., Storz, J.F., Antunes, A., Greenwold, M. J., Meredith, R. W., et al. Comparative genomics reveals insights into avian genome evolution and adaptation. Science, 346(6215):1311–1320, 2014.

22. Zerbino, D. R. and Birney, E. (2008). Velvet: algorithms for de novo short read assembly using de bruijn graphs. Genome research, 18(5), 821–829.

23. Jeffrey A Rosenfeld, Darryl Reeves, Mercer R Brugler, Apurva Narechania, Sabrina Simon, Russell Durrett, Jonathan Foox, Kevin Shianna, Michael C Schatz, Jorge Gandara, et al. Genome assembly and geospatial phylogenomics of the bed bug cimex lectularius. Nature communications, 7, 2016.

24. Shubha Vij, Heiner Kuhl, Inna S Kuznetsova, Aleksey Komissarov, Andrey A Yurchenko, Peter Van Heusden, Siddharth Singh, Natascha M Thevasagayam, Sai Rama Sridatta Prakki, Kathiresan Purushothaman, et al. Chromosomal-level assembly of the asian seabass genome using long sequence reads and multi-layered scaffolding. PLoS Genet, 12(4):e1005954, 2016.

